# Macrophage Migration Inhibitory Factor (MIF) Overexpression Induced by SARS-CoV-2 Infection Promotes Neural Regeneration in Human Brain Organoids

**DOI:** 10.64898/2026.02.13.705720

**Authors:** Andrea Martí-Sarrias, Maria C. Puertas, Isabel Turpin-Moreno, Lidia Garrido-Sanz, Ángel Bayón-Gil, Patricia Resa-Infante, Jakub Chojnacki, Maria C. Garcia-Guerrero, Víctor Urrea, Ramón Lorenzo-Redondo, Sandra Acosta, Javier Martinez-Picado

## Abstract

While SARS-CoV-2 primarily targets the respiratory system, its neurological effects have become a significant clinical concern. Postmortem analyses reveal astrogliosis, neuronal death, and blood-brain barrier dysfunction, yet the interplay between neural injury and endogenous repair remains unclear. Here, we employed human embryonic stem cell–derived brain organoids to examine viral tropism, bystander effects, and regenerative responses following infection. Single-cell transcriptomics and histological assays showed that SARS-CoV-2 productively infects neurons, neural progenitors, astroglia, and choroid plexus cells, triggering widespread apoptosis and senescence in both infected and neighboring cells. Despite low infection rates, organoids activated robust regenerative programs, including axon guidance, Wnt pathway signaling in mature neurons, and radial glia proliferation. Importantly, macrophage migration inhibitory factor (MIF) emerged as a key mediator, being strongly upregulated in both infected and uninfected cells, particularly in the choroid plexus. Recombinant MIF promoted dendritic outgrowth and cortical progenitor activation in uninfected organoids. Computational analyses indicated that MIF stimulates neural regenerative via EGFR signaling and upregulates its own expression in non-infected cells. These findings identify MIF as a molecular link between SARS-CoV-2-induced neural damage and regenerative activation in cortical cells.

## INTRODUCTION

The emergence and rapid global spread of severe acute respiratory syndrome coronavirus 2 (SARS-CoV-2), with over 780 million cases reported cases late 2025 (*COVID-19 Cases* | *WHO COVID-19 Dashboard*, n.d.; Dong et al., 2020), has profoundly affected public health. While SARS-CoV-2 primarily targets the respiratory system, it also infects diverse cell types and tissues, leading to extrapulmonary manifestations. Neurological effects became apparent during the initial pandemic wave associated with the Wuhan variant, prior to widespread vaccination (Havers et al., 2022). During the acute phase, up to 36% of COVID-19 cases exhibited neurological symptoms, with higher rates observed among hospitalized and older individuals (Mao et al., 2020; Singh et al., 2022; Taquet et al., 2021).

Postmortem studies and experimental models have demonstrated that SARS-CoV-2 infection leads to multiple neuropathological changes–including astrogliosis, microgliosis, axonal degeneration, neuronal death, and increased blood-brain barrier (BBB) permeability–even when viral loads in the brain are low (García-González et al., 2023). These findings suggest that neural injury may result from bystander mechanisms, where infected cells impact neighboring tissue. However, molecular insights into these processes are limited, mainly due to the challenges inherent to postmortem analyses. Additionally, neuropathological changes do not always correlate with clinical symptoms, especially in cognitively resilient individuals. Issues such as the lack of appropriated healthy controls, variability in postmortem intervals, differences in pre-analytical procedures, and inconsistent tissue preservation can all compromise sample quality and confound interpretation of results (Birdsill et al., 2011; Danner et al., 2024; Ferrer et al., 2007; Ma et al., 2024). Given these inherent limitations of postmortem analyses, human stem cell-derived brain organoids provide a robust and controlled platform for studying viral neurotropism and pathogenesis, effectively reducing confounding factors. While several studies have shown that SARS-CoV-2 infects brain organoids *in vitro* (Jacob et al., 2020; McMahon et al., 2021; Mesci et al., 2022; Pellegrini et al., 2020; Ramani et al., 2020; Samudyata et al., 2022; Song et al., 2021), the subsequent effects on neural tissue homeostasis and regeneration remain controversial.

While some patients experience prolonged neurological symptoms during acute SARS-CoV-2 infection, over 60% recover fully within approximately seven months (Beretta et al., 2023). Clinical outcomes are highly variable, particularly in younger individuals, who are more often asymptomatic and report neurological symptoms in only 1-20% of cases (Chua et al., 2021). This heterogeneity suggests that intrinsic repair mechanisms may be activated in response to virus-induced brain injury, with neural progenitor cells and neurons potentially contributing to the restoration of neural homeostasis.

Given the complex interplay between neural injury and repair mechanisms observed following SARS-CoV-2 infection, it is essential to examine molecular factors that may bridge inflammation and regeneration within the brain. Macrophage migration inhibitory factor (MIF)stands out as a central molecule at the crossroads of neuroinflammation and tissue repair. This pleiotropic cytokine plays dual roles: it drives immune responses by promoting pro inflammatory mediators such as tumor-necrosis factor (TNF), interferon-γ (IFN-γ), and interleukin-12 (IL-12) during infections with intracellular pathogens (Koebernick et al., 2002), and is upregulated in systemic inflammation (Calandra et al., 2000) and neurodegenerative diseases, like Alzheimer’s (Bacher et al., 2010; Hok-A-Hin et al., 2023). Conversely, MIF also promotes neural stem cell proliferation and neurogenesis *in vitro* and *in vivo* (Bank et al., 2012; Ohta et al., 2012, 2016). However, its function as a molecular link connecting SARS-CoV-2-induced neural injury to subsequent regenerative processes remains largely unexplored.

In this study, we employed human brain organoids to model SARS-CoV-2 infection in the central nervous system (CNS) and investigated the relationship between neural injury and regeneration. Through single-cell transcriptomics and histological analyses, we identified molecular signatures indicative of both direct and bystander neuronal responses to SARS-CoV-2. Notably, we found that MIF is strongly upregulated after viral exposure, serving as a critical mediator of regenerative signaling in neural tissue. These findings reveal new aspects of SARS-CoV-2-related neuropathology and identify MIF as a promising therapeutic target to influence neural recovery after infection.

## RESULTS

### SARS-CoV-2 virus infects, replicates, and spreads in neural cell types

To investigate the neurotropism and replication capacity of SARS-CoV-2 in neural cells, we generated unguided whole-brain organoids from H9 human embryonic stem cells (hESC) and differentiated them for 90 days before infection (**Figure 1A**). Immunofluorescence (IF) analysis confirmed the development of major brain cell types and the establishment of cortical cytoarchitecture. By day 30, organoids contained radial glia (Nestin^+^), cortical progenitors (PAX6^+^), outer radial glia (HOPX^+^), and early born neurons (TUBB3^+^), including a subpopulation of mature deep-layer neurons (MAP2^+^, FOXP2^+^) (**Supplementary figure 1A**). By day 90, mature MAP2^+^ NeuN^+^ neurons had expanded, upper-layer SATB2^+^ neurons had appeared, and progenitors (Nestin, HOPX, PAX6) persisted alongside an emerging astroglia population (GFAP^+^). Additionally, regions of pseudostratified epithelium expressing TTR indicated the presence of mature choroid plexus (ChP) cells (**Figure 1B, Supplementary figure 1B**). Differentiation was terminated at day 90 to ensure the presence of all major neural cell populations required for subsequent infection assays while maintaining temporal efficiency.

**Figure 1.**
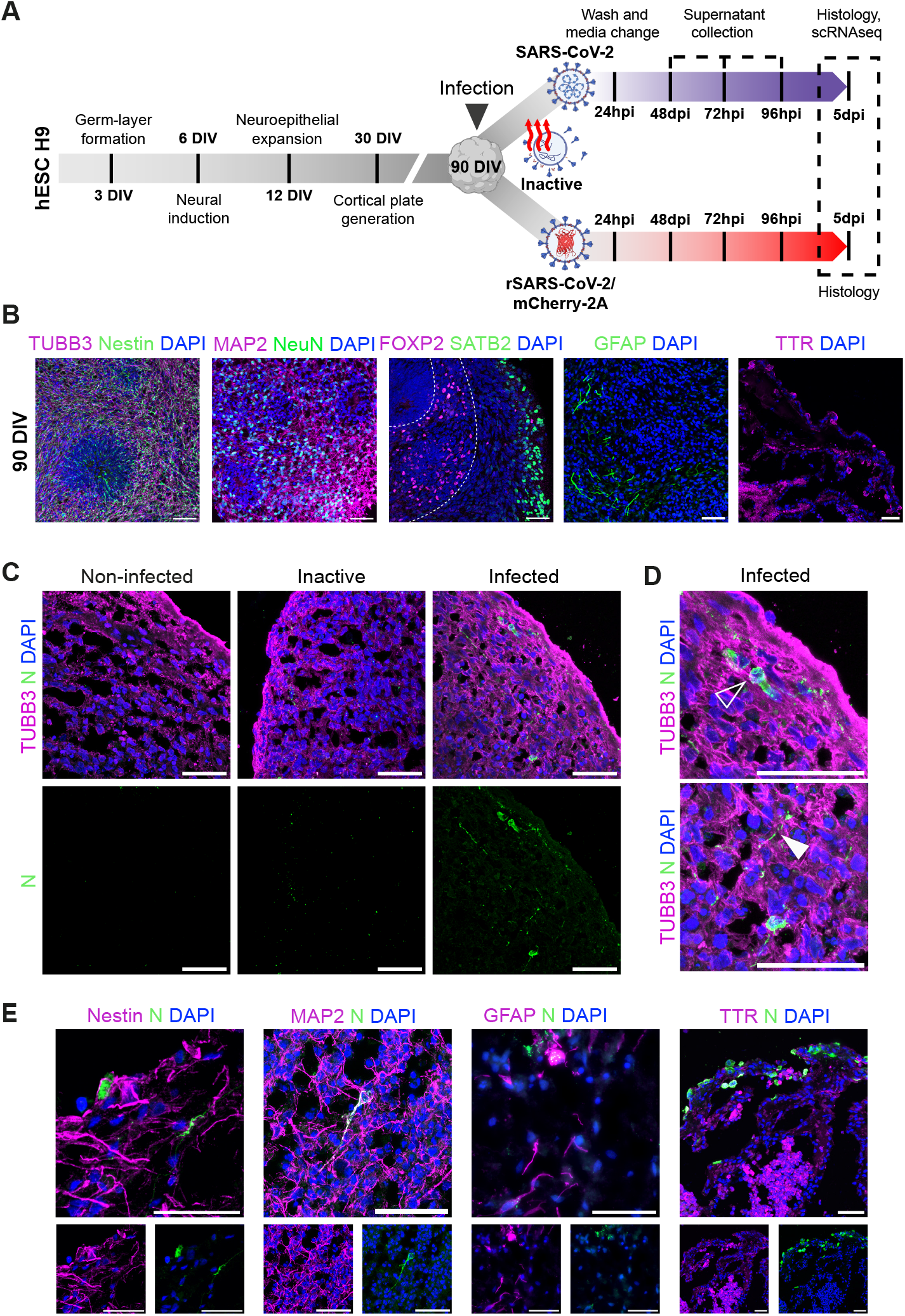
SARS-CoV-2 actively infects human-derived brain organoids. **A.** Schematic representation of the experimental workflow. Human brain organoids were differentiated for 90 days and then exposed to SARS-CoV-2 virus (MOI 0.1), heat-inactivated SARS-CoV-2 virus, or recombinant rSARS-CoV-2/mCherry-2A (MOI 0.1). hESC: human embryonic stem cells. MOI, multiplicity of infection. DIV, days in vitro. hpi, hours post-infection. dpi, days post-infection. **B**. Cell populations in brain organoids at day 90 of differentiation. Immunofluorescent images to visualize markers for neural progenitors (Nestin, green), immature neurons (TUBB3, magenta), mature neurons (MAP2, magenta; NeuN, green), deep-layer neurons (FOXP2, magenta), upper-layer neurons (SATB2, green), astroglia (GFAP, green), and mature choroid plexus (ChP) cells (TTR, magenta). **C**. Immunofluorescence analysis of organoid sections showing SARS-CoV-2 N protein (N, green) in immature neurons (TUBB3, magenta). Nuclei are stained with DAPI (blue). Squares highlight expanded regions. **D**. Inset of highlighted regions from C, including neuronal somas (empty arrowhead) and axons (white arrowhead). **D**. SARS-CoV-2 N protein was also detected in Nestin^+^ neural progenitor cells, MAP2^+^ mature neurons, TTR^+^ ChP cells, and GFAP^+^ astroglia, indicating broad tropism of the virus within the organoid cell population. Scale bars: 50 µm.

To assess whether neural cells in our brain organoid model support SARS-CoV-2 infection and replication, as previously demonstrated in studies using immortalized cell lines (Chu et al., 2020; Zhang et al., 2020), we exposed 90-day-old organoids to the virus for 24 hours and monitored viral replication over five days (**Figure 1A**). Immunofluorescence analysis of viral nucleocapsid (N) protein revealed a sparse yet consistent infection pattern, with infected cells predominantly localized at the organoid periphery (**Figure 1C, Supplementary figure 2A**). Notably, organoids exposed to heat-inactivated virus showed minimal intracellular N protein, supporting that the brighter N staining observed following infection with replication competent-virus was attributable to active infection. Higher magnification imaging confirmed perinuclear and axonal localization of N protein (**Figure 1D**). Co-immunostaining detected viral N protein in neural progenitor cells (Nestin^+^), mature neurons (MAP2^+^), astrocytes (GFAP^+^) and ChP cells (TTR^+^), with ChP cells showing the highest susceptibility to SARS-CoV-2 infection (**Figure 1E)**, consistent with previous reports (Jacob et al., 2020; Pellegrini et al., 2020). Of note, SARS-CoV-2 infection did not induce macroscopic changes in organoids (**Supplementary figure 2A**), in contrast to other neurotropic viruses (Qian et al., 2017).

Viral replication was assessed by quantifying viral RNA levels in culture supernatants at three and five days post infection. Organoids exposed to infectious SARS-CoV-2 exhibited a progressive increase in viral RNA, whereas those exposed to heat-inactivated virus showed no detectable replication across multiple organoid replicates (**Figure 2A**). To confirm viral replication, we used a recombinant SARS-CoV-2 strain encoding the fluorescent reporter mCherry (rSARS-CoV-2/mCherry-2A), which is expressed fused to viral 2A non-structural protein only during active viral replication and not incorporated into viral particles. Infected brain organoids displayed strong colocalization of viral N protein with mCherry in Nestin^+^ neural progenitors, NeuN^+^ mature neurons, GFAP^+^ astrocytes, and TTR^+^ ChP cells (**Figure 2B**). No N or mCherry signal was detected in non-infected controls or organoids exposed to heat inactivated virus (**Supplementary figure 2B, Supplementary figure 2C**). Collectively, these findings demonstrate that SARS-CoV-2 productively replicates in all major neural cell types within human brain organoids, including progenitors, neurons, astroglia, and ChP cells.

**Figure 2.**
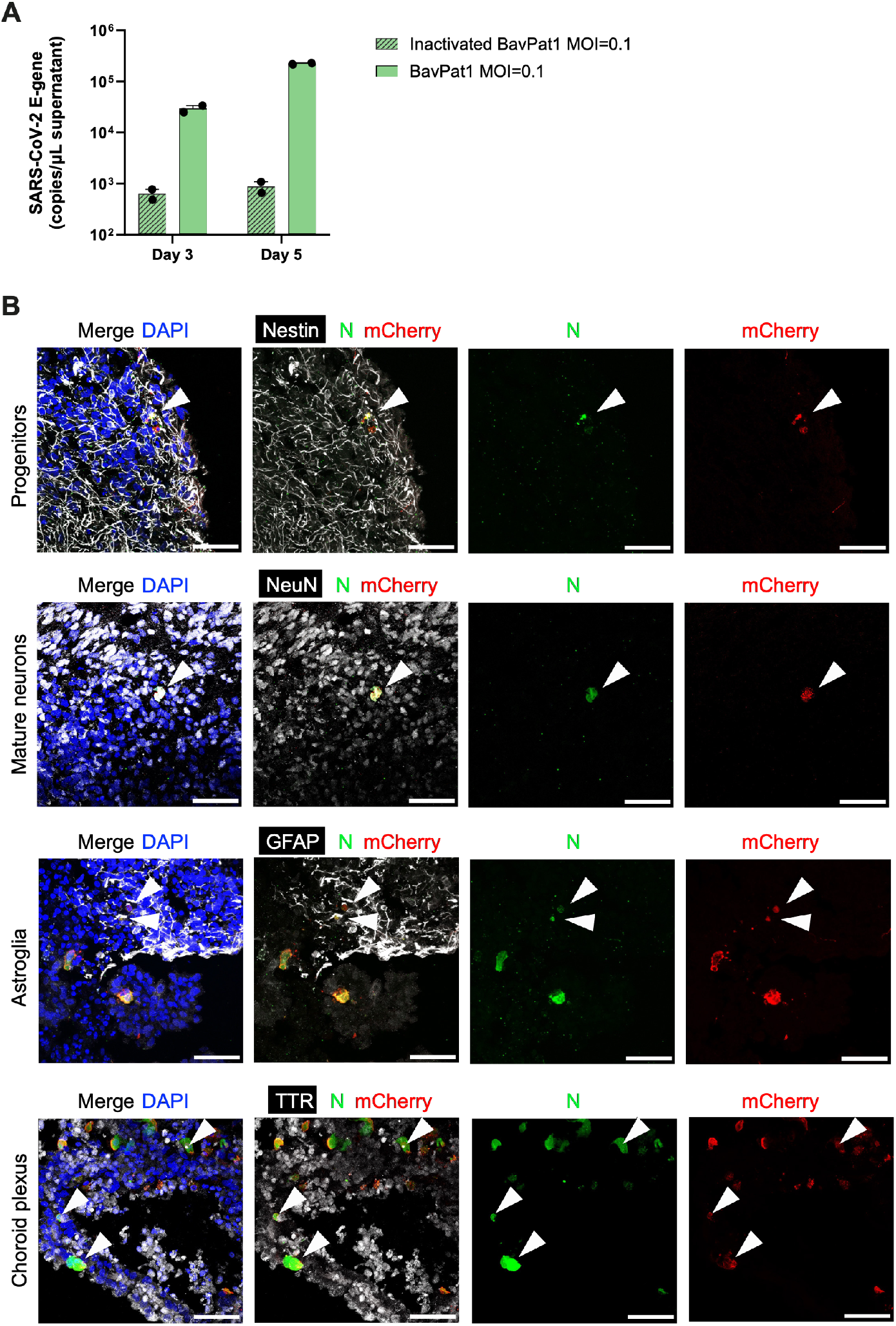
SARS-CoV-2 infects and replicates in human-derived brain organoids. **A.** Quantification of viral E gene copies/µL in the culture supernatant by RT-qPCR, confirming SARS-CoV-2 (BavPat1 strain, MOI 0.1) replication in infected organoids. **B**. Immunofluorescence analysis of brain organoids infected with recombinant rSARS-CoV-2/mCherry-2A (MOI 0.1) reveals co-localization of viral SARS-CoV-2 N protein (N, green) and mCherry reporter (red) in multiple cell types, including neural progenitors (Nestin, white), mature neurons (NeuN, white), astroglia (GFAP, white), and ChP cells (TTR, white). Nuclei are stained with DAPI (blue). White arrowheads indicate double-positive cells (N^+^mCherry^+^), representing productive infection. MOI, multiplicity of infection. Scale bars: 50 µm.

### SARS-CoV-2 infection and replication disrupt cellular homeostasis across neural cell types

To examine the impact of SARS-CoV-2 infection on distinct neural cell populations, we performed single-cell RNA sequencing (scRNA-seq) on infected and uninfected brain organoids five days after viral inoculation. Following rigorous quality control, normalization, dimensionality reduction, and clustering, we generated a unified UMAP embedding that integrated both infected and uninfected samples (**Figure 3A**). Cell clusters were manually annotated based on canonical marker expression profiles observed in uninfected brain organoids (**Figure 3B**).

**Figure 3.**
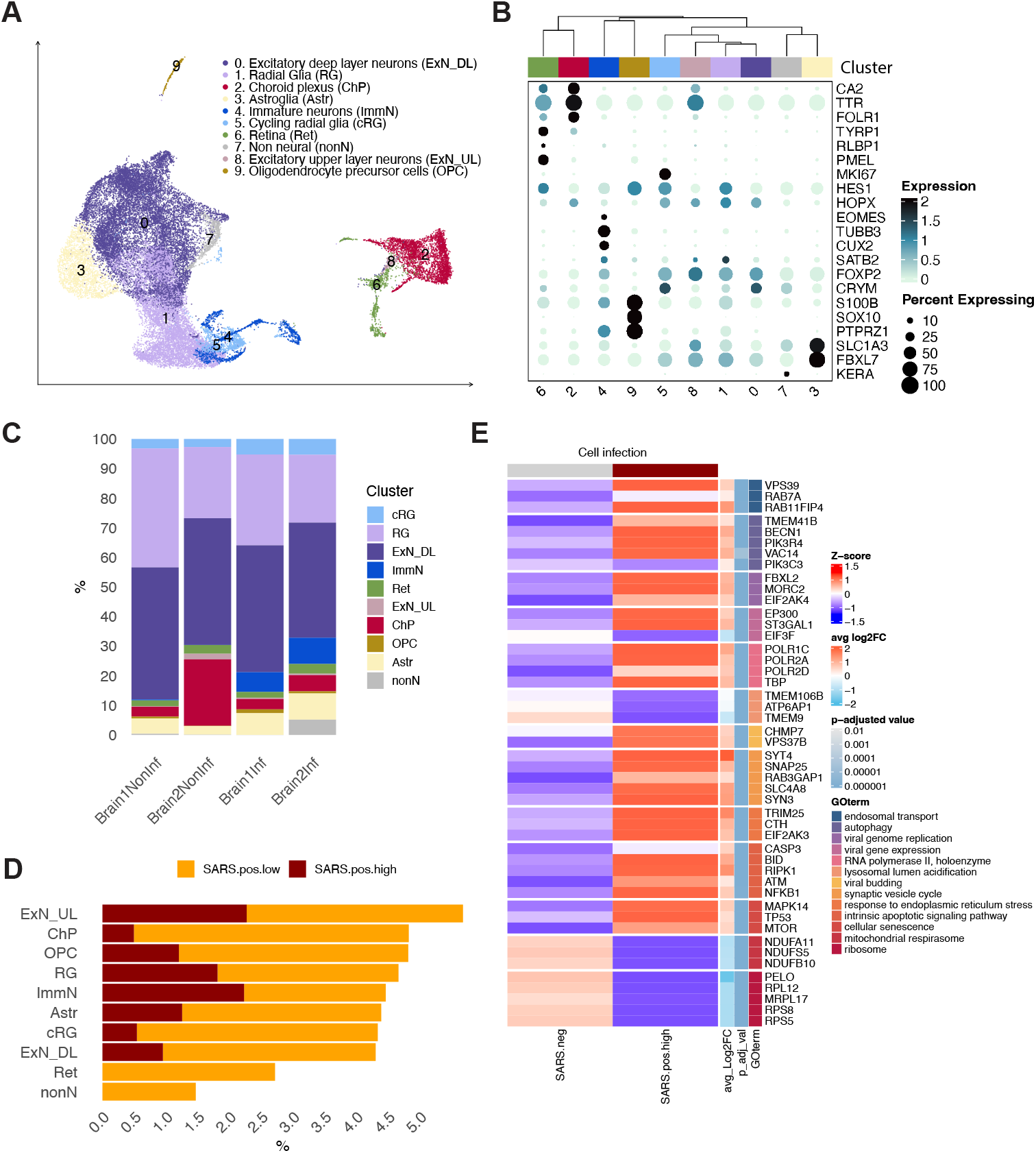
Single-cell RNA sequencing analysis of SARS-CoV-2-infected cells in brain organoids. **A.** UMAP projection showing the clustering of cells from both infected and uninfected brain organoids. Ten distinct clusters were identified: excitatory deep-layer neurons (ExN DL, cluster 0), radial glia (RG, cluster 1), choroid plexus cells (ChP, cluster 2), astroglia (Astr, cluster 3), immature neurons (ImmN, cluster 4), cycling radial glia (cRG, cluster 5), retinal cells (Ret, cluster 6), non-neural cells (nonN, cluster 7), excitatory upper-layer neurons (ExN UL, cluster 8) and oligodendrocyte precursor cells (OPC, cluster 9). **B**. Dotplot showing the expression levels and percentages of marker genes for each cell cluster in uninfected samples. **C**. Percentage of cells per cluster in infected and uninfected organoid samples (two pools of brain organoids, with 3 organoids each). **D**. Quantification of infected cells per cluster based on normalized expression of the viral N gene. Cells were grouped by N viral transcript abundance into three cell infection categories: SARS.neg (N = 0), SARS.pos.low (0 < N ≤ 1) and SARS.pos.high (N > 1). **E**. Heatmap of differentially expressed genes across infection categories (SARS.neg, SARS.pos.high) with their corresponding Gene Ontology (GO) term from the enrichment analysis. Z-scores are calculated based on the average expression values per cell per cell infection category including the SARS.pos.low category. Average Log2FC and p adjusted values refer to the differential gene expression test between SARS.pos.high and SARS.neg cells.

We identified both neural and non-neural cell populations, including: deep-layer excitatory neurons (ExN DL, cluster 0) characterized by robust expression of *FOXP2, CRYM*, and *FOS*; radial glial cells (RG, cluster 1) expressing *HES1, S100B*, and *HOPX*; choroid plexus cells (ChP, cluster 2), positive for *TTR* and *CA2*; astroglia (Astr, cluster 3), marked by *VIM, SLC1A3* (GLAST), and *FBXL7*; and cortical immature neurons (ImmN, cluster 4), predominantly expressing *TUBB3*. Additionally identified populations included cycling radial glia (cRG, cluster 5), characterized by *MKI67* and *HES1* expression; retinal cells (Ret, cluster 6), comprising photoreceptors expressing *TYRP1* and *RLBP1* as well as retinal pigmented epithelium expressing *PMEL*; a small population of upper-layer excitatory neurons (ExN UL, cluster 8), expressing *CUX1, SATB2*, and *HTR2C*; oligodendrocyte precursor cells (OPC, cluster 9), expressing *S100B, SOX10*, and *PTPRZ1*; and a distinct non-neural population (nonN, cluster 7), predominantly found in one infected sample and characterized by expression of *KERA*.

Notably, infected organoids showed an increased proportion of immature neurons and cycling radial glia compared to uninfected controls (**Figure 3C**), indicating that SARS-CoV-2 infection may alter neural cell composition and disrupt cellular homeostasis within the organoid model.

To evaluate viral infection at the transcriptomic level, we assessed the expression of SARS-CoV-2 genes across the identified cell clusters. Among the viral genes, *N* and *ORF10* were the most abundantly detected transcripts. Notably, viral N RNA was present in up to 4.3% of cells, including neurons, radial glia, astroglia, ChP cells, and OPCs (**Figure 3D**). Based on the abundance of N transcripts, cells were classified into three infection stages: high (SARS.pos.high; N > 1), low (SARS.pos.low; 0 < N ≤ 1), and negative (SARS.neg; N = 0). While all cell population clusters contained between 1% and 5% of cells with detectable N viral transcripts, only 0.5% to 2% of cells showed high levels of viral transcripts, indicative of productive infection. This elevated N expression, observed across all major neural populations was consistent with our histological observations (**Figure 1E)**.

We next examined transcriptomic alterations in cells with high levels of SARS-CoV-2 infection (SARS.pos.high) compared to uninfected cells (SARS.neg) (**Figure 3E, Supplementary table 1**). Infected cells displayed significant upregulation of genes involved in endosomal trafficking, including *VPS39* and *RAB7A*, consistent with viral entry via the endosomal pathway (Baggen et al., 2021; Walia et al., 2024). This observation aligns with the observed low expression of *TMPRSS2* and the widespread presence of cathepsins B (*CTSB*) and L (*CTSL*) in the brain, as the endosomal route is favored in cells with little to no *TMPRSS2* expression (Kettunen et al., 2023). Furthermore, SARS CoV-2 infection further triggered the activation of host mechanisms required for viral genome replication (Zhou et al., 2023). Notably, genes associated with autophagy and double-membrane vesicle formation, including *TMEM41B, BECN1*, and *PIK3R4*, were upregulated, consistent with the formation of viral replication complexes. In addition, genes implicated in viral genome replication, such as *EP300*– a histone acetyltransferase relevant for SARS-CoV-2 (Ochsner et al., 2020; Rebendenne et al., 2022; Salgado-Albarrán et al., 2021)–were also upregulated. Increased expression of RNA polymerase II genes further supported enhanced viral transcriptional activity in infected cells (**Figure 3E, Supplementary table 1**).

Additionally, we detected reduced expression of genes responsible for lysosomal lumen acidification, such as *TMEM106B, ATP6AP1* and *TMEM9*, suggesting that SARS-CoV-2 may increase the pH within the ER-Golgi intermediate compartment and lysosomal compartments, thereby promoting an environment conducive to viral particle assembly–a mechanism previously implicated in viral pathogenicity in Vero cells (Wang et al., 2023) (**Figure 3E, Supplementary table 1**). Conversely, we observed increased expression of genes implicated in viral budding, particularly members of the CHMP and VPS protein families that comprise the endosomal sorting complexes required for transport (ESCRT) machinery. This indicates that SARS-CoV-2 likely exploits the ESCRT pathway for viral egress (**Figure 3E, Supplementary table 1**). Furthermore, infected cells also exhibited upregulation of genes associated with vesicle-mediated synaptic transport, such as *SYT4* and *SNAP25* (**Figure 3E, Supplementary table 1**), suggesting that SARS-CoV-2 may facilitate its intracellularly dissemination through vesicular transport within cortical neurons, as has been described in peripheral nerves (Fenrich et al., 2020).

Finally, SARS-CoV-2 infection triggered stress responses across multiple cellular pathways. Infected cells showed increased expression of ER stress markers, such as *TRIM25*. Furthermore, the intrinsic apoptotic signaling and NF κB pathways were dysregulated in infected cells, evidenced by elevated levels of *CASP3, BID* and *NFKB1*, consistent with previous studies of cell death in infected neural cells (Ramani et al., 2020). Genes associated with cellular senescence–including, the cellular arrestors *CDKN1A, CDKN2A* and *MTOR, TP53, MAPK14*–were also upregulated, indicating activation of multiple stress response mechanisms. Mitochondrial dysfunction was observed through downregulation of complex I subunits (NDUF genes), while ribosomal biogenesis and protein translation were impaired, as reflected by reduced expression of 40S and 80S ribosomal proteins (**Figure 3E**). Together, these findings demonstrate that SARS-CoV-2 causes a profound disruption of homeostasis in infected cells.

### SARS-CoV-2 infection induces bystander cell death, senescence and neuronal regeneration

Although only 0.5 and 2% of cells exhibited productive SARS-CoV-2 infection, we investigated whether infection triggered broader pathological effects in the neuronal population (**Supplementary table 2**). Focused on ExN DL–the most abundant and mature neuronal subtype in our brain organoids–transcriptomic profiling revealed widespread upregulation of apoptotic and stress-related genes, including *FAS, CASP4, DKK1, MIF*, and *IL18*, in non-infected ExN DL neurons from infected organoids compared to controls (**Figure 4A**). Elevated *CASP4* expression suggested that cell death was mediated by extrinsic apoptotic signals. Signature analysis for genes involved in positive regulation of the extrinsic apoptotic signaling pathway (GO:2001238) revealed upregulation in infected organoids, notably, among non-infected (SARS.neg) cells, indicating a bystander effect (**Figure 4B**). Histological quantification using the active cleaved Caspase-3 (aCASP3) marker confirmed increased apoptosis in both infected and neighboring non infected cells (**Figure 4C-D**). Additionally, expression of senescence-associated cyclin-dependent kinase inhibitors *CDKN1A* (p21) and *CDKN2D* (p19) was elevated (**Figure 4A**), and activation of the cellular senescence signature (GO:0090398) (**Figure 4B**) along with increased β-galactosidase (βGal) staining in uninfected neurons (**Figure 4C**), confirmed enhanced neuronal senescence following SARS-CoV-2 exposure. No changes in the MAP2 dendritic marker area were observed at the histological level (**Figure 4-D**).

**Figure 4.**
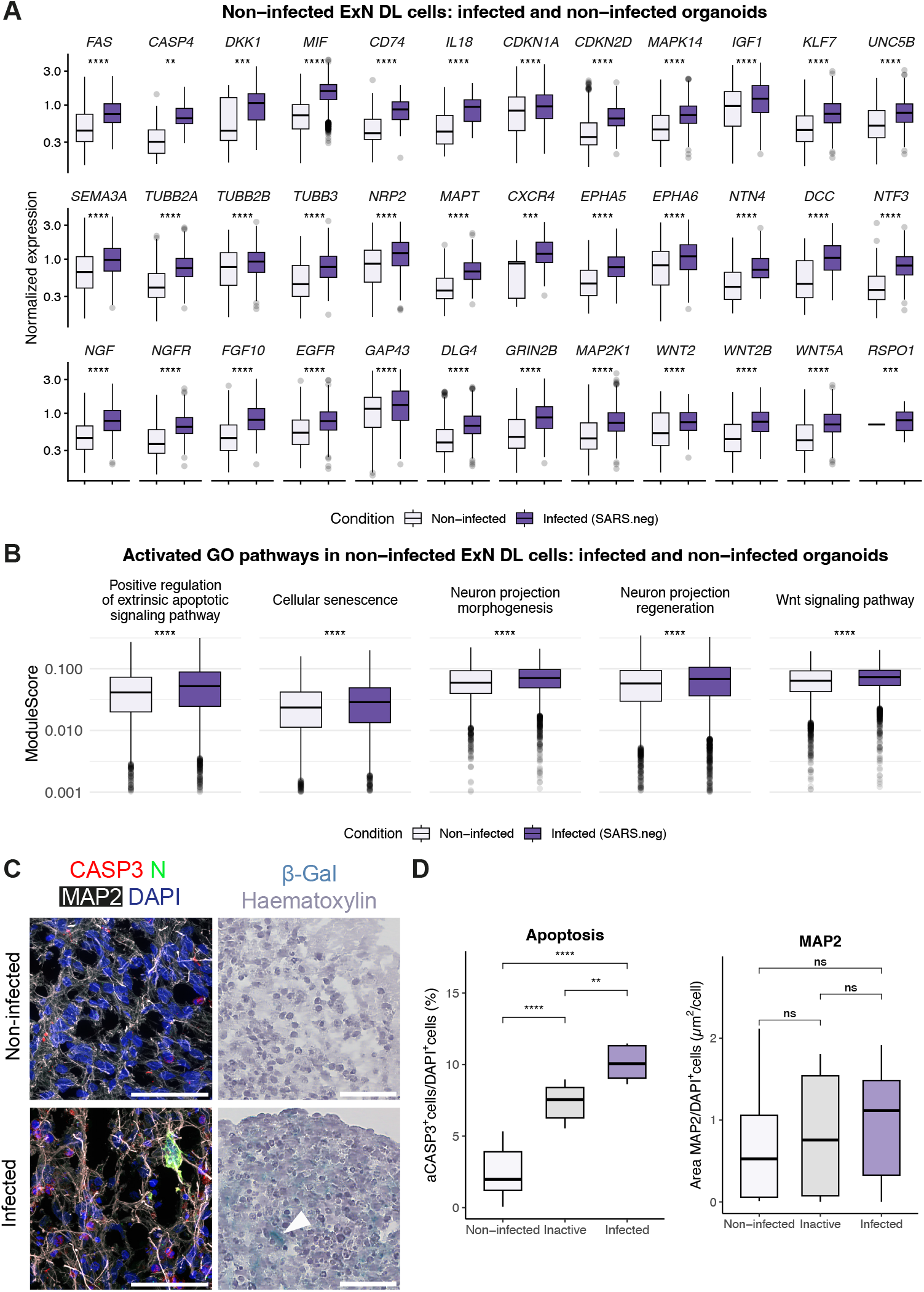
SARS-CoV-2 induces bystander stress and apoptosis in non-infected mature neurons of infected brain organoids. **A.** Upregulated genes with significant p-adjusted value in excitatory deep-layer neurons (ExN DL, cluster 0) from infected vs. non-infected organoids. Normalized expression values are separated according to the infection status of the organoids (infected, non-infected). Only SARS.neg cells are shown. Y axis is represented in log10 scale. Statistical significance determined by unpaired Wilcoxon test with Bonferroni adjustment. **B**. Signatures (module scores) calculated per cell of different Gene Ontology (GO) terms: Positive regulation of extrinsic apoptotic signaling pathway (GO: GO:2001238), cellular senescence (GO:0090398), neuron projection morphogenesis (GO:0048812), neuron projection regeneration (GO:0031102), and Wnt signaling pathway (GO:0016055). Signatures of SARS.neg ExN DL cells from infected and non-infected conditions were compared. Statistical significance determined by unpaired Wilcoxon test; *p<0.05, **p<0.01, ***p<0.001, ****p<0.0001. **C**. Immunofluorescence staining of organoid sections 5 days post-infection showing activated Caspase-3 (aCASP3, red) and SARS-CoV-2 N protein (N, green) in mature neurons (MAP2, white), as well as β-Galactosidase (β-Gal, blue, white arrowhead) staining indicating cellular senescence. Nuclei are stained with DAPI (blue) or Hematoxylin (purple). **D**. Percentage of aCASP3^+^ cells normalized to the total number of cells (DAPI^+^) comparing the non-infected condition, organoids exposed to inactive virus, and infected organoids. A significant increase in apoptotic cells is observed in the inactive and infected conditions. Percentage of MAP2^+^ area relative to the total number of cells (DAPI^+^) indicates no significant change in total dendritic area in the different conditions. For IF quantification, at least 3 pictures from 2 organoids from 2 independent infections were used. Statistical significance determined by unpaired t-test; *p<0.05, **p<0.01, ***p<0.001, ****p<0.0001. Scale bars: 50 µm.

Interestingly, despite these degenerative changes, we also detected transcriptional signatures of neural regeneration and tissue repair in non-infected ExN DL neurons. Genes associated with neuron projection morphogenesis and regeneration–including *CXCR4, DCC, EGFR, EPHA5, EPHA6, FGF10, IGF1, KLF7, MAPT, NGF, NGFR, NRP2, NTF3, NTN4, SEMA3A, TUBB2A, TUBB3, UNC5B, DLG4, GAP43* and *GRIN2B–*were significantly upregulated in SARS.neg neurons from infected organoids. Key components of the MAPK pathway, such as *MAP2K1*, which supports neurite outgrowth and neuron survival (Markus et al., 2002), were also elevated (**Figure 4A**). Signatures for neuron projection morphogenesis (GO:0048812), neuron projection regeneration (GO:0031102), and the Wnt signaling pathway (GO:0016055), including effectors *WNT2, WNT2B, WNT5A*, and the regulator *RSPO1*, were activated (**Figure 4B**), highlighting pathways critical for neurodevelopment and regeneration (Rosso & Inestrosa, 2013). Altogether, these findings indicate that SARS-CoV-2 infection in brain organoids triggers bystander neuronal damage in non-infected cells through multiple pathways while concurrently activating transcriptional programs linked to neural regenerative responses.

### SARS-CoV-2-induced damage stimulates radial glia proliferation and activates cortical neurogenic programs

Prompted by evidence of increased neural growth factor activation, we examined the transcriptional dynamics of neural progenitor populations, focusing on radial glia (RG) and cycling radial glia (cRG), the most immature cell types in brain organoids. Differential gene expression analysis between infected and uninfected samples revealed that neural progenitors in infected brain organoids significantly upregulated genes associated with mitotic progression and DNA replication (*CENPF, MKI67, E2F1*) (**Figure 5A, Supplementary table 2**). These transcriptomic changes were supported by increased numbers of phospho-histone H3 (pH3^+^) cells, indicating heightened proliferation, as shown by tissue immunostaining (**Figure 5B-C**). In addition, genes linked to neurogenesis and neuronal differentiation (*PAX6, LHX2, HES1, HES5, TUBB3, STMN2, NRXN1* and *NCAM1)* were upregulated in RG cells from infected samples, suggesting these cells are primed for neuronal differentiation (**Figure 5A, Supplementary table 2**). Immunofluorescence confirmed an expansion of Nestin-positive regions, consistent with an enlarged progenitor pool (**Figure 5B-C**). These molecular and histological changes coincided with increased frequencies of cRG and immature neuron (ImmN) populations in infected organoids, as observed in scRNAseq analysis (**Figure 3C**). Collectively, these results suggest that SARS-CoV-2 infection induces both proliferative expansion and neurogenic activation in RG cells, likely representing a compensatory regenerative response to local cellular stress and injury within the brain organoid environment.

**Figure 5.**
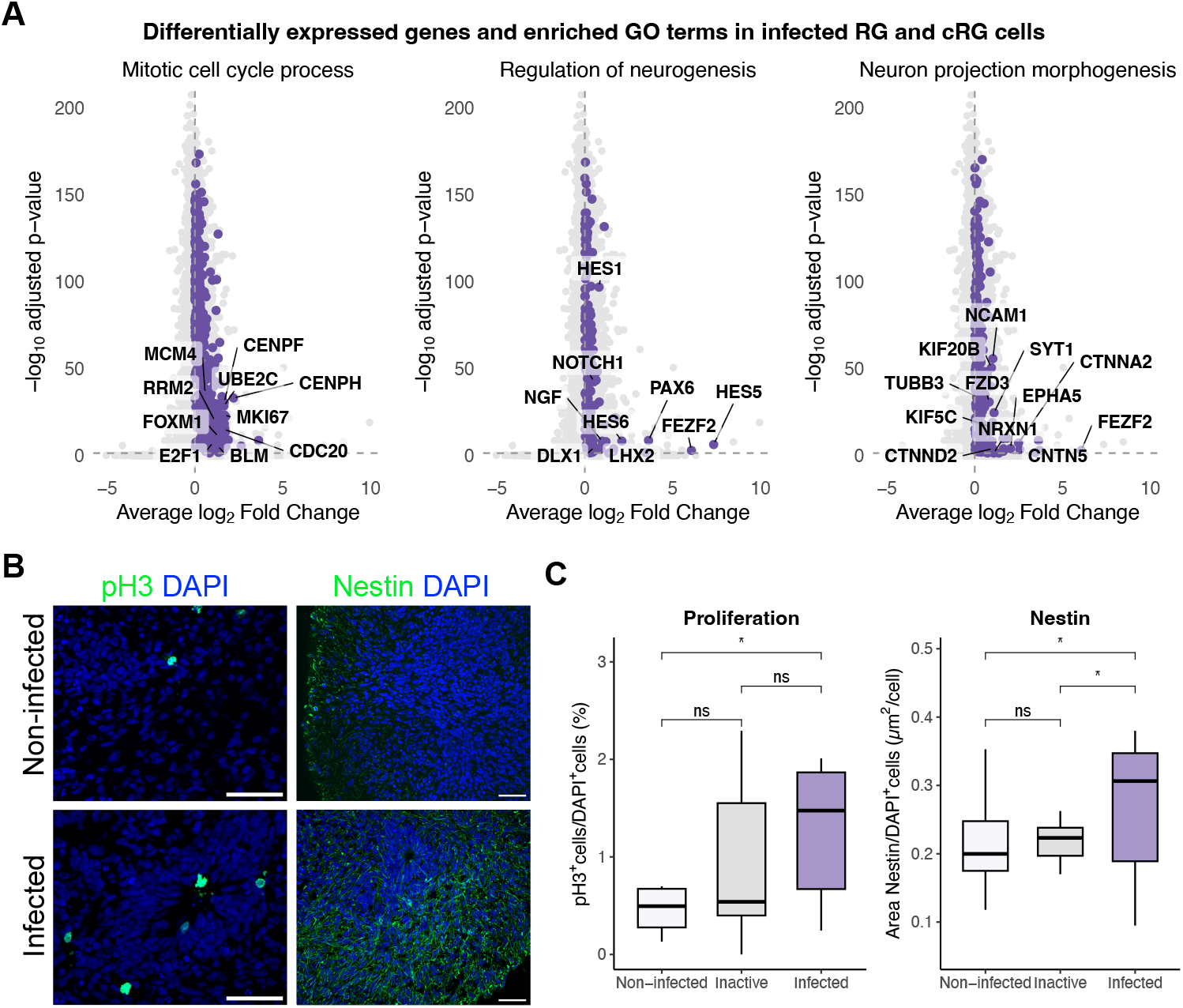
SARS-CoV-2 infection induces proliferation of radial glial cells in brain organoids. **A.** Volcano plot showing differentially expressed genes in radial glial cells (RG and cRG, cluster 1 and 5) in infected vs. non-infected conditions. Purple dots correspond to genes of the enriched Gene Ontology (GO) term: Mitotic cell cycle process (GO:1903047), regulation of neurogenesis (GO:0050767), and neuron projection morphogenesis (GO:0048812). Y axis is log10 scaled. Statistical significance determined by unpaired Wilcoxon test, with Bonferroni adjustment; *p<0.05, **p<0.01, ***p<0.001, ****p<0.0001. **B**. Immunofluorescence staining of organoid sections 5 days post infection showing phospho-histone H3 (pH3, green) and Nestin (green), markers of proliferation and neural progenitors, in non-infected and infected organoids. Nuclei are stained with DAPI. **C**. Percentage of the number of pH3^+^ cells and ratio of the Nestin^+^ area normalized to the total number of cells (DAPI^+^) comparing the non-infected condition, organoids exposed to inactive virus, and infected organoids. A significant increase in the number of proliferating cells and neural progenitor areas is observed following infection. For IF quantification, at least 3 pictures from 2 organoids from 2 independent infections were used. Statistical significance determined by unpaired t-test; *p<0.05, **p<0.01, ***p<0.001, ****p<0.0001. Scale bars: 50 µm.

### Ubiquitous MIF overexpression drives regenerative response in infected organoids

The regenerative signatures observed in non-infected deep-layer excitatory neurons (ExN DL) after SARS-CoV-2 exposure prompted investigation into their molecular drivers. Among candidate mediators **(Figure 4A)**, and based on its described functions, MIF emerged as a potential key inflammation-induced factor linking cellular injury to neuronal repair. While basal *MIF* expression was present in most cells of uninfected organoids, infection led to robust *MIF* upregulation across all cell types with average log_2_ fold changes ranging from 2.56 in ExN DL (non-infected=0.36; infected=1.23) to 3.98 in retina (non-infected=0.51; infected=2.56) and ChP cells (non-infected=0.38; infected=2.50) (**Figure 6A**). Of note, ChP and retina showed the highest *MIF* overexpression, despite scarcity of infected cells in those clusters. Further analysis revealed that in RG and ExN DL–the predominant populations in infected organoids–*MIF* upregulation occurred mainly in non-infected cells (**Figure 6B**). This pattern was consistent with metanalysis of COVID-19 frontal cortex samples, which also showed increased *MIF* expression comparable to that seen in our brain organoid model (**Supplementary figure 3A**) (Mavrikaki et al., 2022). Immunofluorescence confirmed elevated MIF protein levels in infected brain organoids compared to controls, especially in the pseudostratified ChP epithelium (**Figure 6C**). MIF protein localized to both infected and non-infected cells (**Figure 6C**), and quantification of stained area showed a trend towards higher MIF levels in infected samples, further supporting the transcriptomic findings (**Figure 6D**).

**Figure 6.**
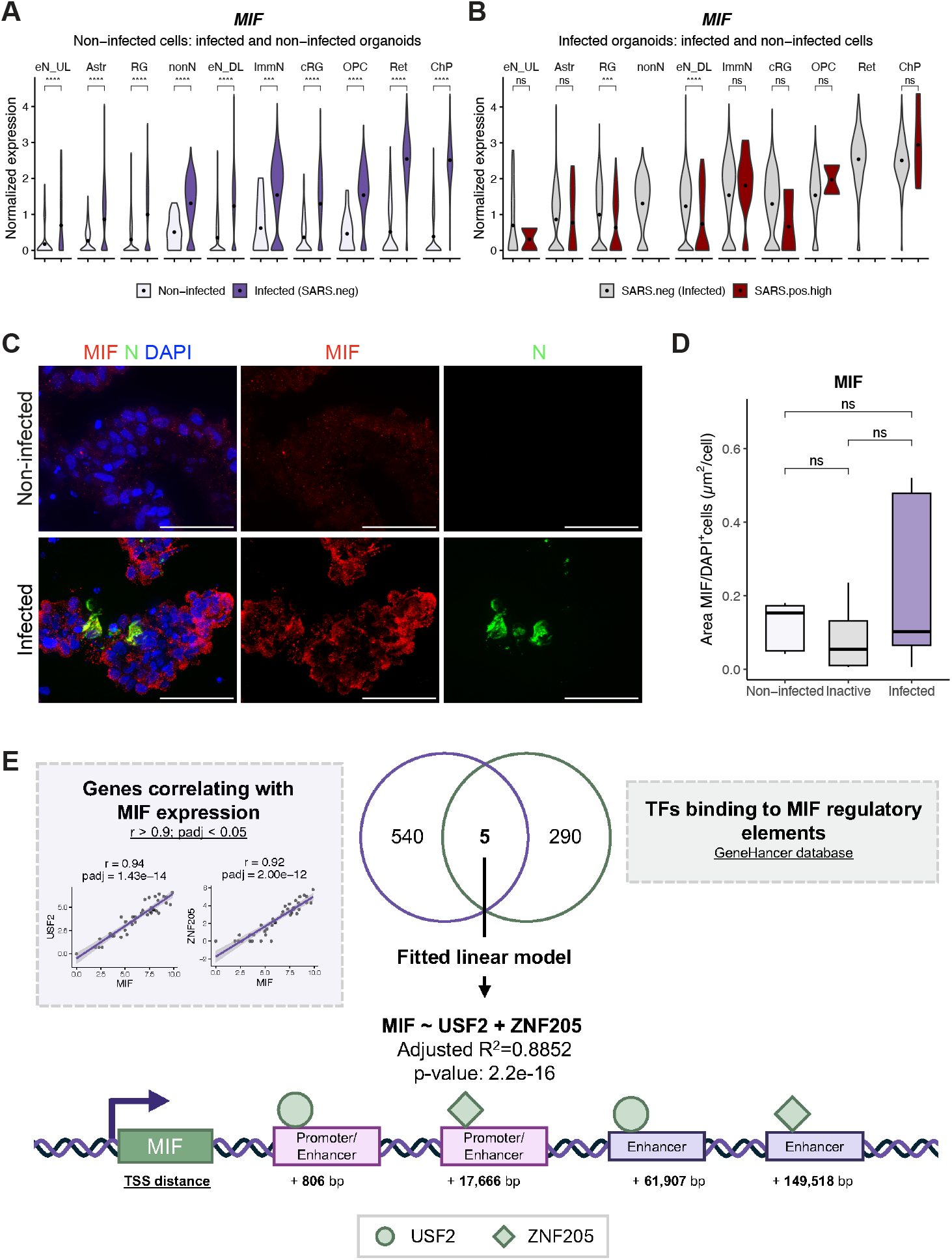
SARS-CoV-2 infection induces upregulation of MIF in brain organoids. **A.** Normalized expression of *MIF* across cell clusters in non-infected and infected organoids, including only SARS.neg cells. Clusters are ordered from left to right according to the increasing difference in mean *MIF* expression between infected and non-infected organoids. **B**. *MIF* expression in infected brain organoids by cluster stratified by viral load: infected (SARS.pos.high) and non-infected cells (SARS.neg). Cell cluster order follows panel A. Dots inside violin plots indicate mean values. Statistical significance determined by unpaired Wilcoxon test: *p<0.05, **p<0.01, ***p<0.001, ****p<0.0001. Absence of brackets indicates no test was performed due to lack of data. **C**. Representative immunofluorescence images of organoid sections 5 days post-infection showing MIF (red) and SARS-CoV-2 N protein (green) in non infected and infected organoid sections. Nuclei are stained with DAPI. **D**. Percentage of MIF^+^ area normalized to DAPI, comparing the non-infected condition, organoids exposed to inactive virus, and infected organoids. For IF quantification, at least 3 pictures from 2 organoids were used. Statistical significance determined by unpaired Wilcoxon test: *p<0.05, **p<0.01, ***p<0.001, ****p<0.0001. Scale bars: 50 µm. **E. Left dotted box**: Pearson coefficients (r) and p adjusted values (p.adj) calculated from log-normalized pseudobulk expression values of *MIF* and the rest of genes in ExN DL cluster. Only positive and significant correlations (r > 0, p.adj < 0.05) were retained. **Right dotted box**: Source used to define candidate TFs binding to promoters or enhancers of *MIF*. **Center**: Venn diagram showing the overlap between genes that positively correlated with *MIF* in ExN DL and genes annotated by GeneHancer as putative regulators of *MIF*. Three genes were common to both sets: HMG20B, ZNF205 and USF2. The combination of *HMG20B, ZNF205* and *USF2* expression could explain 71% of the *MIF* expression variability across all cell classes. **Bottom**: genomic distribution of the identified regulatory elements including promoters and enhancers of MIF. Predicted TFs binding sites based on GeneHancer data are represented with different symbols.

To explore the regulatory mechanisms driving the widespread upregulation of MIF in infected organoids, we identified genes correlating with MIF. Correlation was computed using log-normalized pseudobulk gene expression values across all cell types, samples, and conditions, to reduce single-cell variability, ensure independence and normality between values. Using this method, we identified 543 genes significantly correlated with MIF. Using the GeneHancer database, we found 295 transcription factors (TFs) predicted to bind 34 known proximal and distal *MIF* regulatory elements (**Supplementary table 3**). Of these, five *MIF* TFs–*USF2, ZNF205, ZNF324, MLLT1* and *HMG20B –*were consistently correlated with *MIF* expression across all cell types (**Figure 6E**).

We first fitted a multivariable linear model including all five TFs ordered by Pearson’s coefficient in a descendent manner (linear model: MIF ∼ USF2 + ZNF205 + ZNF324 *+* MLLT1 + HMG20B). We confirmed by ANOVA test that only USF2 and ZNF205 were adding new significant information to the model. Model reduction revealed that USF2 and ZNF205 alone accounted for approximately 88% of the variance in MIF expression (reduced linear model: MIF ∼ USF2 + ZNF205; **Figure 6E**), with *ZNF324, MLLT1* and *HMG20B* contributing to marginal additional explanatory power. Together, these results suggest that USF2 and ZNF205 act as dominant, non–cell-type-specific upstream regulators of MIF, consistent with its broad induction across infected organoids.

To determine cell type-specific upstream regulators of MIF, we performed single-cell correlation analysis within the major clusters: ChP, cRG, RG, ExN DL and ExN UL. Distinct sets of *MIF*-correlated TFs were identified in each cluster, indicating unique regulatory mechanisms driving *MIF* upregulation (**Supplementary figure 3B**). Notably, *MIF* expression positively correlated with the non-cell-type-specific TFs (**Figure 6E**), immediate stress response genes (*FOS, JUN, JUND)* (Zhou et al., 2007), interferon signaling regulators *IRF2–*an antiviral mediator relevant to COVID-19 pathogenesis (Oh et al., 2024)–, retinoic acid receptor pathway members *NR2F1, NR2F2, RARA* and *RXRB* in the ExN DL cluster, which are involved in neuronal progenitor differentiation to mature neurons (Yu et al., 2012), and genes that code for proteins that are known to interact with SARS-CoV-2 viral proteins (Zhou et al., 2022) (**Supplementary figure 3B**). Altogether, these results indicate that rapid *MIF* upregulation following SARS-CoV-2 infection acts as a neural damage response and may contribute to neural regeneration.

To assess the functional consequences of MIF upregulation, and recognizing MIF as a paracrine secreted mediator, we treated non-infected brain organoids with human recombinant MIF protein (1 µg/mL) for 6 days (**Figure 7A**). MIF exposure led to a significant increase in dendritic area, measured by MAP2^+^ immunostaining normalized to DAPI^+^ nuclei. Enhanced dendritogenesis was confirmed by co-staining for MAP2 and NeuN, though it was not accompanied by a rise in neurogenesis (NeuN/DAPI). We also detected elevated levels of the immature neuron marker TUBB3 (**Figure 7B-C**), consistent with effects seen after SARS-CoV-2 infection (**Figure 3C**). Markers for proliferation (pH3) and apoptosis (aCASP3) remained unchanged, indicating that MIF specially promotes dendritic arborization without affecting cell proliferation or death. At the transcriptomic level, *SOX2* did not show significant upregulation, but *WNT3A* expression increased by qPCR following MIF exposure, suggesting MIF may direct neural progenitors towards a cortical fate (**Figure 7D**).

**Figure 7.**
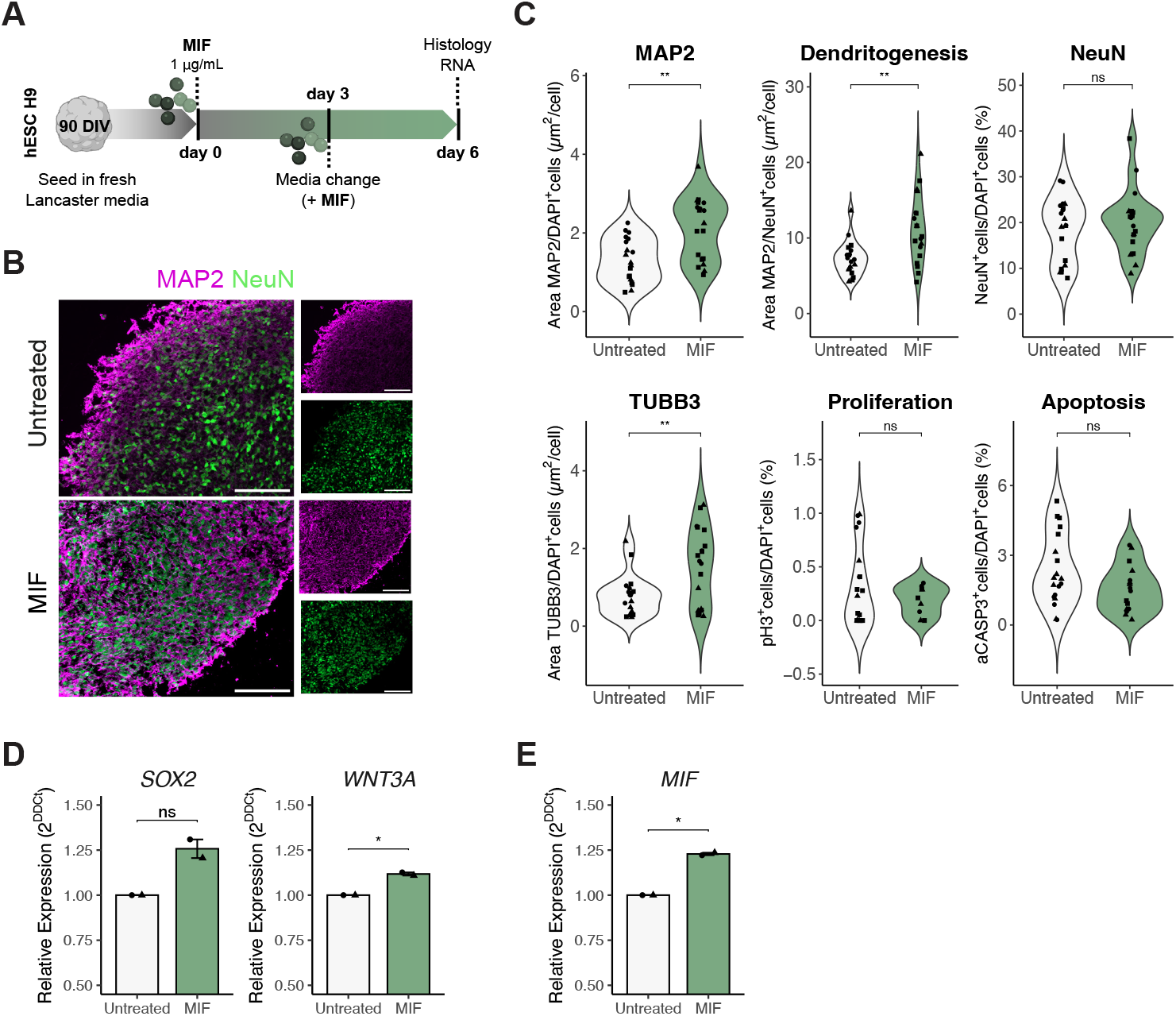
Exposure to exogenous MIF induces dendritogenesis and immature neuron growth in brain organoids. **A.** Schematic of the experimental MIF exposure setup. **B**. Representative immunofluorescence images of MIF-treated organoids showing neural dendrites (MAP2, magenta) and mature neuron nuclei (NeuN, green). **C**. Quantification of immunofluorescent markers after MIF exposure, including neural dendrites (MAP2), dendritogenesis (MAP2/NeuN), mature neuron nuclei (NeuN), immature neurons (TUBB3), proliferation (pH3), and apoptosis (aCASP3). Dot shapes correspond to experimental replicates. Statistical significance determined by unpaired Wilcoxon test: *p<0.05, **p<0.01, ***p<0.001, ****p<0.0001. Scale bars: 50 µm. **D, E**. Relative expression (2^DDCt^) respect to the Untreated condition of *SOX2, WNT3A* and *MIF* after MIF exposure by qPCR. Dot shapes correspond to experimental replicates. Statistical significance determined by unpaired Wilcoxon test: *p<0.05, **p<0.01, ***p<0.001, ****p<0.0001.

To investigate how MIF promotes neuronal regeneration in non-infected cells following SARS-CoV-2 infection, we applied LIANA+ for cell-cell communication inference to our scRNAseq dataset. LIANA+ infers the downstream causal network linking known ligand-receptor interactions with intracellular signaling pathways and TF activation in the receiver cells. This analysis revealed that, in ExN DL neurons (cluster 0), MIF likely signals through EGF receptor (EGFR), with this interaction markedly upregulated after infection (**Supplementary figure 4A**). In these neurons, MIF appears to activate the PI3K/MAPK pathway via EGFR, converging on TFs *MEF2A, SRF, MYF5, MYOD1* and *CTNNB1* (β-Catenin) (**Supplementary figure 4B, Supplementary table 5**), most of which were significantly upregulated post-infection in the ExN DL cluster (**Supplementary table 2**).

Given the robust upregulation of *MIF* in SARS-CoV-2-infected brain organoids—including non-infected bystander cells—we hypothesized that MIF may reinforce its own expression via a paracrine positive feedback loop. Leveraging LIANA+ and predicted MIF-binding TFs (**Supplementary table 3**), we assembled an intracellular signaling network linking MIF signaling to downstream TFs with potential to activate *MIF* transcription, supporting the existence of a self-amplifying loop (**Supplementary figure 5, Supplementary table 4**). The autoregulatory positive feedback was confirmed *in vitro*, as exogenous MIF exposure further increased MIF gene expression in brain organoids **(Figure 7E**).

Taken together, these findings demonstrate that SARS-CoV-2 infection is sufficient to trigger widespread *MIF* expression in brain organoids, where MIF may function as both a neuroprotective and regenerative mediator by promoting dendritic outgrowth, highlighting its dual role in neural damage response and repair.

## DISCUSSION

Although hospitalizations and mortality have declined since the first pandemic wave due to public health interventions, the long-term neurological sequelae of COVID-19 remain a major concern. Understanding the balance between neural injury and repair is crucial. Brain organoids provide a powerful, human-specific model to study neural development, SARS-CoV-2 neurotropism, and neuropathology (Jacob et al., 2020; McMahon et al., 2021; Mesci et al., 2022; Pellegrini et al., 2020; Ramani et al., 2020; Samudyata et al., 2022; Song et al., 2021). Here, we combined single-cell transcriptomics and histological analyses to dissect the cellular consequences of SARS-CoV-2 infection in human brain organoids, shedding light on cell susceptibility, viral entry and replication mechanisms, and the downstream neural responses–including both injury and regeneration–elicited even by low-level infection in the brain.

We confirmed that SARS-CoV-2 infects multiple neural cell types, including RG, ExN DL, astroglia, OPCs, and ChP cells, with viral replication observed across neuronal, glial, and epithelial lineages. This supports the notion of a broad cellular tropism of the virus in the brain. Mechanistically, infection relies primarily on the endosomal entry pathway, as indicated by low *TMPRSS2* expression and the upregulation of cathepsin-dependent endocytosis, aligning with previous findings in neural and epithelial systems (Hoffmann et al., 2020; Ou et al., 2020).

Despite the low frequency of infected cells in our brain organoids—mirroring the scarce detection of SARS-CoV-2 in post-mortem brain tissues from COVID-19 cases (Matschke et al., 2020)—we observed widespread transcriptional changes indicative of cellular stress, including activation of apoptotic and senescence pathways, in both infected and uninfected cells. This suggests bystander effects that propagate cellular dysfunction throughout neural tissue, consistent with observations in other viral infections (Hippee et al., 2021; Martínez-Mármol et al., 2023; Mesci et al., 2022; Song et al., 2023, 2021). Increased apoptosis and senescence in non-infected regions were confirmed by activated cleaved Caspase-3 and β-Gal staining, highlighting a neurotoxic environment after infection. Interestingly, we also detected upregulation of *CASP4* in ExN DL, marking inflammatory programmed cell death (pyroptosis), which is characterized by membrane rupture (Broz, 2025; Liu et al., 2024), and has been described in other viral infections affecting the brain such as rabies virus, Zika, and herpesvirus (He et al., 2020; Hu et al., 2022; Yu et al., 2024), and in monocytes during acute COVID-19 (Ferreira et al., 2021).

Degenerative alterations were accompanied by regenerative transcriptional signatures, notably upregulation of neural projection molecules and Wnt pathway components, particularly in ExN DL neurons. This dual response suggests a compensatory mechanism aiming to restore tissue integrity after viral injury. The inherent plasticity of developing brain tissue may partially explain the lower incidence of neurological sequelae in pediatric COVID-19 (Chua et al., 2021). Among neuronal repair pathways, MIF emerged as a key modulatory factor. *MIF* expression was elevated throughout infected organoids, particularly in non-infected cells as part of the bystander effect and was most abundant in ChP cells. MIF, previously linked to immune regulation and neural injury, has context-dependent pro-inflammatory and neuroprotective roles (Leng et al., 2003; Furukawa et al., 2017). Its upregulation has been reported in cortical tissue from COVID-19 donors (Mavrikaki et al., 2022; Zhang et al., 2024) and circulating soluble MIF levels positively correlate with COVID-19 disease severity (Aksakal et al., 2021), especially in individuals carrying high-expression genetic variants (Shin et al., 2023). In Alzheimer’s disease, MIF is found upregulated (Hok-A-Hin et al., 2023), and may defend neurons against Aβ-induced toxicity (Zhang et al., 2019). Infected ChP cells are the main MIF source upon infection, while uninfected bystander cells may activate *MIF* expression via a positive feedback mechanism. MIF secretion by ChP cells, along with other proteins, to the cerebrospinal fluid (CSF) has been linked to enhanced neurogenesis and neurite outgrowth in mouse embryos (Courtney et al., 2025). Additionally, *MIF* transcription can also be activated through Toll-like receptors (TLRs), facilitating bystander induced expression (Yao et al., 2016).

Our analysis revealed that *MIF* upregulation across multiple neural and non-neuronal cell types is regulated by both shared and cell type-specific TFs. *HMG20B, ZNF205*, and *USF2* explained much of the *MIF* transcriptional variance, suggesting they act as global regulators in neural tissue. HMG20B is a chromatin regulator of neuronal differentiation and synaptic maturation (Maddhesiya et al., 2025; Wynder et al., 2005). USF2 binds the *BDNF* promoter in neurons and may influence development and synaptic plasticity (Chen et al., 2003), while ZNF205, less characterized in the CNS, is part of a protein family implicated in several neurological disorders (Bu et al., 2021). At the cell type-specific level, immediate early-response genes (*FOS, JUN*, and *JUND)*, and *IRF2* (an antiviral and interferon response mediator) positively correlated with *MIF* expression in deep-layer excitatory neurons. Consistent with MIF’s pleiotropic role in tissue remodeling, inflammation, and regeneration, MIF exposure to neural cultures induced neurite outgrowth and primed progenitors towards a cortical fate (Courtney et al., 2025; Zhang et al., 2013).

Cell-to-cell communication analysis further revealed that MIF downstream effectors involved in neuronal repair (MEF2A, SRP, CTNNB1) act via EGRF and PI3K/MAPK signaling. These TFs are known to regulate neurogenesis, synaptogenesis, neurite remodeling, and regeneration: β-Catenin activates the Wnt pathway for cortical neurogenesis (Wang et al., 2022), MEF2A promotes mature synaptic bouton formation (Yamada et al., 2013), and SRF regulates dendritic growth and spine maturation (del Blanco et al., 2019). Robust *MIF* upregulation in uninfected cells is likely mediated by a positive feedback mechanism, where soluble MIF enhances its own production.

In summary, SARS-CoV-2 infection in human neural tissue triggers a complex response involving direct viral replication, secondary bystander injury, activation of apoptotic and senescence pathways, and engagement of regenerative programs. MIF emerges as a central molecular node connecting infection-driven damage with reparative signaling, offering new insights into host responses and potential therapeutic targets. Collectively, this work underscores the importance of investigating both viral neuroinvasion mechanisms and endogenous regenerative processes that shape disease progression and recovery in the CNS.

## METHODS

### Human embryonic stem cell (hESC) culture and brain organoid generation

The human embryonic stem cell (hESC) line WA009 H9 was maintained in mTeSR1 (85850, Stem Cell Technologies) and passaged using Gentle Cell Dissociation Reagent (100-0485, Stem Cell Technologies) when they were at ∼70% confluency. For BO generation, hESC colonies at ∼80% confluency were detached using Gentle Cell Dissociation Reagent, pipetted to have a single-cell suspension, and resuspended in mTeSR1 supplemented with ROCK Inhibitor Y-27632 (RI) 10 nM (HY-10583-10, Medchem). For non-guided brain organoid protocol, 9000 cells per well were seeded in a U-bottom ultra-low attachment 96-well-plate (CLS7007-24EA, Corning). When cells formed aggregates with smooth edges, media was changed to mTeSR1 without RI. When aggregates reached 500 μm in diameter, typically by day 3 after seeding, media was changed to Neural Induction Media containing DMEM-F12 (11330057, Gibco) supplemented with 1% N2 (A1370701, Gibco), 1% GlutaMAX (35050061, Gibco), 1% MEM-NEAA (11140050, Gibco) and 1ug/mL heparin. Between days 4–6, when edges became bright, distinctive that primitive neuroepithelium was appearing, brain organoids were embedded in Matrigel droplets using 8 μL Matrigel (45354277, Cultek) per organoid, and transferred to 50 mm non-adherent petri dishes (124-17, Termofisher) in Differentiation Media without Vitamin A (IDM−A) consisting of 48% DMEM-F12, 48% Neurobasal (21103049, Gibco), 1% GlutaMAX, 1% penicillin–streptomycin (P/S; 15070063, Gibco), 0.5% MEM-NEAA, 0.5% N2, 1% B27 supplement without vitamin A (12587010, Gibco), 0.035% 1:100 2-mercaptoethanol and 25 μg/mL insulin (I9278-5ML, Sigma). After 7 days in IDM-A, media was changed to Differentiation Media with Vitamin A (IDM+A) and organoids are transferred to a 19-mm orbital shaker set to 62 rpm. Half media was changed twice a week.

### SARS-CoV-2 viral stocks

Human 2019-nCoV reference viral isolate (strain BavPat1/2020) was produced at Charité - Universitätsmedizin Berlin (CUB) and distributed via EVAg network. The viral stock was further expanded and titrated in Vero E6 cells (ATCC, CRL-1586) using a limiting dilution assay, with infectious unit concentrations calculated by the Reed– Muench method (Scotti, 2013), and the preservation of the furin cleavage site was verified by sequencing. The recombinant rSARS-CoV-2/mCherry-2A, kindly provided by L.Martinez-Sobrido, was generated from an artificial bacterial chromosome containing the cDNA infectious clone of SARS-CoV-2 USA-WA1/2020 strain (Chiem et al., 2021). Viral stock was generated by transfection of the pBAC with X-tremeGENE 9 DNA Transfection Reagent and further titrated in VeroE6 cultures.

### *In vitro* SARS-CoV-2 infection

Three-months old brain organoids were exposed to SARS-CoV-2 virus MOI 0.1 for 24 hours. As non-replicative control, an aliquot of SARS-CoV-2 virus was inactivated by heating at 56ºC for 30 min. To stop the exposure to the virus, brain organoids from each well were washed with 5ml culture media and transferred to new culture plates with fresh IDM+A media. Culture supernatants were collected at day three and five after infection to monitor virus replication dynamics, and brain organoids were harvested 5 days after infection for histological and scRNA-seq analysis.

### Quantification of viral RNA in culture supernatants

QIAamp Viral RNA Mini Kit (Qiagen), E_Assay First Line Screening (IDT, 10006804) and TaqManTM Fast Virus 1-Step Master Mix (Applied Biosystems, 4444434) were used to quantify the number of SARS-CoV-2 genomes by qPCR on the 7500 RealTime System (Applied Biosystems) as previously described (Corman et al., 2020). Serial dilutions of the plasmid 2019-nCoV_E Positive Control (IDT, 10006896) were used as standard curve.

### Recombinant MIF exposure in brain organoids

Brain organoids at 90 days in vitro (DIV) were treated with human recombinant MIF protein (300-69, PeproTech) for 6 days at µg/mL. Half media change was performed on day 3, with fresh recombinant MIF added. Experiments were performed in 3 independently differentiated batches, with 4 brain organoids each.

### Gene expression analysis by quantitative PCR

To evaluate gene expression by qPCR, samples were resuspended in RLT buffer (74104, Qiagen). Total RNA was extracted using the RNeasy Mini Kit (74104, Qiagen) following the manufacturer’s protocol. cDNA synthesis was performed using M-MLV Reverse Transcriptase (M1302, Merck) following the manufacturer’s protocol. qPCR was conducted using SYBR Green Master Mix (A25742, Life Technologies) using ROX passive reference dye. Primer sequences are included in **Table 1**. Expression levels were normalized to *ACTB* housekeeping gene and relative gene expression (2^DDCt^) was calculated by comparison with the non-treated condition.

**Table 1.**
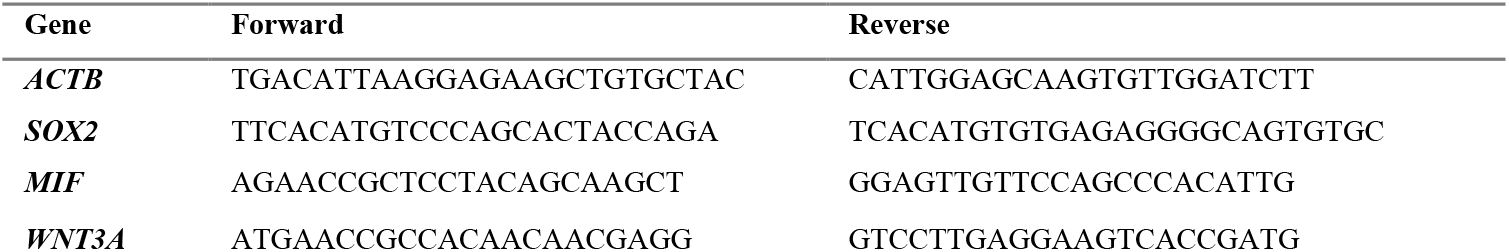
Primers for qPCR.

### Immunostaining

Organoids were fixed in 4% paraformaldehyde (P6148-1KG, Merck) for 45 minutes at RT in agitation, cryoprotected in 30% sucrose overnight at 4ºC, frozen in O.C.T. (29801N, ap medical) using dry ice, and cryosectioned at 12μm. For immunostaining,sections were blocked using 3% donkey serum (D9663, Merck), 2% BSA (A2153, Merck) and 0.1% Triton-X (X100, Merck) for 1 hour. For detection of SATB2 and FOXP2, sections were permeabilized for 10’ with SDS 1% prior to the blocking step. Sections were incubated with primary and secondaries antibodies diluted in blocking buffer as indicated in **Table 2**. Washes were done using PBS-T 0.1%. Sections were mounted using Fluoromount-G (00-4958-02, Thermo).

**Table 2.** Antibodies used for immunostainings.

| Antibodies | Ref | Company | Description |
| --- | --- | --- | --- |
| Anti-TUBB3 [Tuj1 clone], mouse | 801201 | BioLegend | 1:1000, 4°C O/N |
| Anti-MAP2, chicken | ab5392 | Abcam | 1:1000, 4°C O/N |
| Anti-NeuN, rabbit | ab177487 | Abcam | 1:500, 4°C O/N |
| Anti-GFAP, chicken | AB5541 | Merck | 1:1000, 4°C O/N |
| Anti-GFAP, rabbit | GTX108711 | GeneTex | 1:500, 4°C O/N |
| Anti-NESTIN, rabbit | 841901 | BioLegend | 1:500, 4°C O/N |
| Anti-PAX6, rabbit | GTX113241 | GeneTex | 1:500, 4°C O/N |
| Anti-Activated Caspase 3, rabbit | C8487 | Sigma | 1:2000, 1h30' RT |
| Anti-SARS-CoV-2 Nucleocapsid (N) [HL455- MS clone], mouse | GTX635712 | GeneTex | 1:500, 4°C O/N |
| Anti-mCherry, chicken | NBP2-25158 | Novus | 1:500, 4°C O/N |
| Anti-SATB2, rabbit | NBP1-92365 | Novus | 1:500, 4°C O/N |
| Anti-FOXP2 [CL5312 clone], mouse | NBP2-61413 | Novus | 1:200, 4°C O/N |
| Anti-MIF, rabbit | A22682 | ABclonal | 1:200, 4°C O/N |
| Anti-Rabbit AlexaFluor 488, donkey | A-21206 | Thermo | 1:1000, 1h30' RT |
| Anti-Mouse AlexaFluor 594, donkey | 715-585-150 | Jackson ImmunoResearch | 1:500, 1h30' RT |
| Anti-Chicken Alexa Fluor 647, donkey | 703-605-155 | Jackson ImmunoResearch | 1:1000, 1h30' RT |
| DAPI | D1306 | Invitrogen | 1:10000, 5' RT |

### Immunostaining analysis and quantification

IF images for quantification were obtained with 20x/0.8NA dry objective using Carl Zeiss Axio Imager M2 Apotome microscope. Maximum intensity projection was obtained using “Orthogonal projection” from the Zeiss software. Staining quantifications were performed using the open-source software ImageJ (Schindelin et al., 2012). For immunofluorescence quantification, 3-4 pictures were used per organoid, 2-3 organoids per condition, from at least 2 independent experiments. For staining quantification, we generated a mask by setting a threshold based on the histogram fluorescent intensity. Thresholds were set using available algorithms in ImageJ (Table 3). Quantifications for DAPI and aCASP3 were obtained from 20x images using “Watershed” segmentation algorithm followed by “Analyze Particles” tool with following parameters: size=15-Infinity and circularity=0.10-1.00.

**Table 3.** ImageJ threshold algorithms for immunofluorescence quantification.

| Staining | ImageJ threshold algorithm |
| --- | --- |
| DAPI | Default |
| MAP2 | Otsu / Li |
| NeuN | Triangle |
| NESTIN | Otsu |
| CASP3 | Triangle |
| MIF | Moments |
| **TUBB3** | Default |
| **DCX** | MaxEntropy |
| **SOX2** | Tiangle |
| **GFAP** | MaxEntropy |

For counting the number of positive cells for corresponding marker, the intersection between the staining mask and the DAPI mask was calculated with the “AND” option from “Image Calculator”. For boxplot representations and statistical tests, values below Q1 or above Q3 were considered outliers and therefore removed from the analysis.

### Single-cell RNA sequencing

Two pools of brain organoids, with 3 organoids each, were infected with SARS-CoV-2 at an MOI=0.1 as described above. Two additional pools of organoids were left uninfected as negative control. After 5 days of infection, each sample pool was dissociated with 400µL TrypLE™ Express (12605010, Termofisher) supplemented with 100U/mL DNase (LK003170, Worthington) and 10 µM ROCK Inhibitor (Y-27632, StemCell Technologies) for 1 hour at 37ºC. After collection of cells in suspension and further mechanical disruption, 1 volume of the dissociation mix was added and incubated at 37ºC until no cell aggregates could be observed. Enzymatic digestion was blocked with 10% FBS. Then, cell debris was removed using a 40µm filter unit (EasyStrainer, 542040, Greiner Bio-one) and a cell suspension was prepared at 1000 cells/µL in PBS containing 0.05% albumin for GEM generation, barcoding and cDNA library construction using the Chromium Next GEM Single Cell 3ʹ Kit v3.1 and the Chromium Controller system (10x Genomics), following the manufacturer’s protocol.

We concatenated the human and SARS-CoV-2 genomes to generate a hybrid reference for bioinformatic analysis. Thus, GRCh38 genome and its annotation were obtained from 10x Genomics repository (refdata-gex-GRCh38-2020-A.tar.gz), and SARS-CoV-2 assembly ASM985889v3 was downloaded from NCBI and its annotation from EMBL repository (https://covid-19.ensembl.org/info/website/index.html).

FASTQ files were quality control assessed with FASTQC v0.11.9 (Andrews, 2010) and aligned against the hybrid reference using count subcommand from 10x Genomics Cell Ranger v7.1.0 (Zheng et al., 2017). Filtered count matrices were analyzed with R (version 4.3.0) (R Core Team, 2014) on a stable environment created with renv package (version 0.17.3) (Ushey & Wickham, 2025). To retain only good quality individual cells, we used Seurat (version 5.0.1) (Hao et al., 2023) to remove cells having fewer than 600 features, more than 7500 features, and a mitochondrial percentage greater than 20%. Doublets were detected and eliminated using scDblFinder package (version 1.14.0) (Germain et al., 2022). On the RNA assay, we normalized, looked for variable genes, and scaled the feature counts from matrices. We also used raw counts from RNA assay to generate a SCT assay with SCTransform function and mitochondrial and ribosomal percentages were included as variables to regress. Individual samples were integrated with Harmony method (Korsunsky et al., 2019). We processed the SCT assay using a linear dimensional reduction with Principal Component Analysis (PCA), and a non-linear dimensional reduction with Uniform Manifold Approximation and Projection (UMAP). After computing the nearest-neighbor graph construction with default Euclidean distance and dims argument set to 1:25, cells groups into clusters using the default Louvain algorithm and a resolution of 0.1. Clusters were manually annotated using cell-type specific markers from the literature. In samples with SARS-CoV-2 infected organoids, we identified infected cells using the N gene and classified them as “SARS negative” when N expression was below 0, “SARS positive low” when N expression ranged between 0 and 1, and “SARS positive high” for cells with an expression of N greater than 1.

We looked for differentially expressed genes (DEGs) using FindMarkers function with min.pct=0.1 and logfc.threshold parameter set to -Inf, MAST method, and using samples’ ID (orig.ident) as latent variables. Using this function and parameters we compared conditions per cluster (infected *versus* non-infected brain organoids), and infection status within infected samples (SARS positive high *versus* SARS negative). For enrichment analysis, we only kept DEG genes with an adjusted p-value below 0.05, and separated them according to the average log2FC sign (upregulated: avg_log2FC > 0; downregulated avg_log2FC < 0). Enrichment analysis was implemented using the enrichGO function from the ClusterProfiler package (version 4.8.1) (Wu et al., 2021). For cross-dataset validation, we downloaded the supplementary tables from Mavrikaki et al., which included differential expressed genes obtained from RNA-seq analysis from post-mortem cortical brain samples of acute COVID-19 patients and matched ICU/ventilated controls. MIF expression changes from these data were compared with the ones obtained from our dataset, ensuring that both analyses used the same p-value adjustment method.

### Identification of upstream regulators of MIF

To identify general upstream regulators of MIF we generated log-normalized summed counts (“pseudobulk”) for each identity class using the AggregateExpression function from Seurat. Identity classes were based on Samples’ ID (orig.ident), condition (infected, non-infected) and Seurat clusters (0-9). Pseudobulk expression matrix was normalized with log() function from R. The normalized pseudobulk matrix was used to check for genes whose expression correlated with *MIF* by calculating the Pearson correlation estimate (r) per gene using the cor.test function. Bonferroni correction was applied to correct for multiple testing. Correlation was considered significant when p-adjusted values were below 0.05 and r above 0.9. Genes were then intersected with genes suggested to bind *MIF* regulatory elements (enhancers and promoters) from GeneHancer database (**Supplementary table 4**). GeneHancer database is based on regulatory elements from 9 sources: RefSeq, VISTA Enhancer Browser enhancers, EPDnew promoters, UCNEbase ultra-conserved noncoding elements, FANTOM5, Ensembl, ENCODE, dbSUPER, and CraniofacialAtlas (Fishilevich et al., 2017). The linear model was generated using lm() from stats R package, with MIF expression as the response variable and the five intersected transcriptional regulators as predictors in descendent order of r (linear model: MIF ∼ USF2 + ZNF205 + ZNF324 + MLLT1 + HMG20B). Then, Type I ANOVA was used to identify the predictors that were significantly contributing to additional explanatory power. Only significant predictors were considered (reduced linear model: MIF ∼ USF2 + ZNF205). To identify cell-specific upstream regulators of MIF, we applied the same procedure but using log-normalized data matrices per cell in the different Seurat clusters.

### Cell–cell communication (CCC) inference with LIANA+

To infer the intracellular signaling pathways linking extracellular MIF signaling to downstream transcriptional programs in receiver cells, we applied LIANA+ to our scRNA-seq dataset. LIANA+ uses ligand–receptor (LR) interactions as inputs and TFs as outputs to identify the most plausible molecular intermediates connecting both elements (Dimitrov et al., 2024). We used LIANA+ to connect extracellular MIF signaling to downstream transcriptional programs in the receiver ExN DL (cluster 0) population. For the analysis, normalized Seurat matrix was converted to an AnnData object. To establish the “inputs” of the network, we first performed a LR inference using the complete DEG list from the FindMarkers function between the different conditions by clusters, and used the li.multi.df_to_lr function from LIANA. Only ligands and receptors expressed in at least 10% of the cells (expr_prop=0.1) were retained.

To focus on the effect of MIF, we subset the LR table to ExN DL targets and aggregated interaction log2FCs by receptor, retaining all the receptor candidates as network “inputs”. For the candidate TF “outputs”, we estimated TF activities from the DEG list per condition using CollecTRI regulons with decoupler/ULM and selected the top 15 TFs in the ExN DL cluster as outputs. We then downloaded a signed, directed protein–protein interaction (PPI) prior from OmniPath (keeping edges with consensus direction, stimulation/inhibition sign, and curation_effort ≥5) and generated a knowledge graph for the ExN DL cluster with li.mt.build_prior_network() computing node weights as the fraction of cells expressing each gene, to favor broadly expressed intermediates. We then used CORNETO/CARNIVAL to solve the shortest sign-consistent path in the network that explains the obtained MIF-driven receptors (inputs) to the selected TFs (outputs). For this, we used the li.mt.find_causalnet() function and set a penalization in nodes that are expressed in less than 10% of cells of the receiver cluster (node_cutoff=0.1; min/max node penalty=0.01/1.0). To find the shortest path, we also included a penalization score to the addition of new edges to the network (edge_penalty=0.1).

For the positive feedback loop analysis of MIF, we repeated the LIANA+ pipeline using the same network “inputs” but changing the candidate TF “outputs” to the putative regulatory elements of *MIF*. The same parameters were used.

## Acknowledgements

This project has been supported by MSD (3D4COVID) and Fundació La Marató de TV3 (grant 202130-30-31-32, Spain) to the laboratories of JM-P and SA. Research in JM-P’s laboratory is also supported in its immunovirology research by the Spanish Ministry of Science, Innovation and Universities (grants PID2022-139271OB-I00 and CB21/13/ 00063, Spain), and the UNDINE project, which has received funding under the Horizon Europe Research and Innovation programme (grant agreement No 101057100). However, the views and opinions expressed do not necessarily reflect those of the European Union or the granting authority European Union’s Horizon Europe research and innovation program. Neither the European Union nor the granting authority can be held responsible for them. Work at IrsiCaixa is supported by 2021 SGR 00452 and CERCA programme by Generalitat de Catalunya. Research in SA’s laboratory is partially funded by (PID2021-128028-NB-I). This publication was supported by the European Virus Archive GLOBAL (EVA-GLOBAL) project that has received funding from the European Union’s Horizon 2020 research and innovation programme under grant agreement No 871029.

Fellowships, SA is a Serra-Hunter fellow of AGAUR (Generalitat de Catalunya). Lidia Garrido-Sanz received financial support from the grant SLT02823 000257, funded by the “Strategic plan for research and innovation in health” (PERIS), from the Catalan Department of Health.

